# Pathway-resolved flux decomposition reveals hidden kinetic hierarchy in protein folding

**DOI:** 10.64898/2026.05.20.726702

**Authors:** Betül Uralcan, Manuel Alejandro López, Amir Haji-Akbari

**Author notes:** B.U. and M.A.L. contributed equally to this work.

## Abstract

Proteins fold through ensembles of competing pathways, yet the kinetic contribution of each route remains difficult to quantify. Structure-prediction methods such as AlphaFold identify folded endpoints, but do not resolve folding kinetics, pathway heterogeneity, or how flux partitions among competing mechanisms. Here, we introduce a framework that directly decomposes folding flux into pathway-specific kinetic contributions by combining forward-flux sampling with trajectory-level unsupervised learning, avoiding millisecond-scale trajectories, biasing potentials, and *a priori* state discretization. Applied to 2,637 statistically representative folding events of the TC5b variant of Trp-cage, the framework recovers a folding time in near-quantitative agreement with experiments and identifies four pathways distinguished by the ordering of helix formation, hydrophobic collapse, and salt-bridge stabilization. The resulting decomposition shows that structural prevalence is a poor proxy for kinetic importance: the most populated pathways are not the fastest, whereas a rare helix– salt-bridge route is disproportionately efficient and a premature salt bridge produces a frustrated slow route. By assigning statistical weights to competing pathways, this framework links structural evolution to kinetic relevance in biomolecular rare events and reveals how folding landscapes select kinetically important routes from many plausible structural sequences.

## I. INTRODUCTION

Proteins are central to biology, yet how they reach their native functional structures remains incompletely understood. A central unresolved challenge is to determine not only which folding pathways are available, but also how much each contributes to the overall folding kinetics. It has long been established that a protein’s amino-acid sequence encodes its three-dimensional folded structure [1]. While recent advances such as AlphaFold enable reasonably accurate prediction of native structures [2, 3], they capture only endpoints of folding– static minima on rugged energy landscapes [4]. In contrast, folding kinetics emerges from an ensemble of competing pathways, whose individual rate contributions determine both mechanism and outcome. Quantifying such rates is essential for understanding how rapidly proteins acquire function *in vivo* and how mutations, solution conditions, or crowding can redirect folding toward non-productive states. However, no existing framework directly decomposes the total folding flux into pathway-specific rate contributions with statistically meaningful, well-defined weights from atomistic simulations. Consequently, the relative kinetic importance of competing pathways remains largely inferred rather than directly computed. Resolving this limitation is essential for connecting structural evolution to kinetics and for understanding how competing molecular routes govern biomolecular transitions more broadly, including processes such as binding and conformational switching.

Existing experimental and computational approaches provide important but incomplete perspectives on this problem. Bulk and single-molecule experiments can estimate overall folding rates and, in some cases, identify intermediates [5–11], but lack the spatiotemporal resolution needed to reconstruct atomistic mechanisms or quantify pathway-specific kinetics [12]. The ensuing scarcity of detailed kinetic data hampers AI-based efforts to learn folding pathways and timescales. Molecular dynamics (MD) simulations offer a complementary route but have their own limitations. Long unbiased trajectories typically yield too few statistically independent transitions to resolve kinetic weights of competing pathways [13–16]. Enhanced-sampling methods efficiently characterize equilibrium ensembles and free-energy landscapes, but do not directly provide kinetics [17–22], which must instead be inferred through *post hoc* approaches such as transition state theory [23], infrequent meta-dynamics [24], or trajectories from putative transition states [25]. The attained estimates are often sensitive to collective variables and are difficult to validate. Markov state models (MSMs) partially address these limitations by assembling kinetic networks from short trajectories [26–28], but the resulting pathways and rates are sensitive to state discretization, lag times, and the assumption of Markovian dynamics [29, 30]. Weightedensemble approaches [31, 32] can also reconstruct folding rates [33, 34] from ensembles of short trajectories, but pathway-specific rate decomposition generally requires additional structural classification and flux bookkeeping, particularly for distinctions orthogonal to the chosen collective variable. As a result, even for extensively studied fast-folding systems [15], a direct and quantitative decomposition of folding flux into pathway-specific rate contributions remains lacking.

A prototypical example is Trp-cage [35], a 20-residue miniprotein that folds on the microsecond timescale and exhibits key features of larger globular proteins. In its native state, Trp-cage (Fig. 1a) adopts a compact structure in which an N-terminal *α*-helix packs against a Cterminal motif containing a 3_10_-helix and a polyproline-II character to enclose the central Trp side chain. Despite extensive experimental [5, 8, 22, 35–38] and computational [13–18, 20–22, 38–48] studies, its folding mechanism remains debated, with competing collapse-first [40, 41] and helix-first scenarios [38, 44–46], as well as heterogeneous pathways involving transient mispacked intermediates [17, 43]. Path sampling approaches, such as transition path/interface sampling [49, 50], generate reactive trajectories, but in practice, pathway populations inferred from finite path-sampling ensembles can remain sensitive to initialization and undersampling of low-flux routes [41, 42, 47]. More generally, initializing path ensembles from a limited number of time-reversed unfolding trajectories may restrict the sampled folding ensemble toward the chosen starting paths, especially when folding sequences are not simple reversals of unfolding. MSMs have revealed additional intermediates, yet typically rely on *a priori* state discretization and Markovian assumptions [48]. As a result, although prior studies have provided valuable mechanistic insights, they do not yield a direct and statistically robust decomposition of folding flux into pathway-specific rate contributions. The existence of multiple plausible folding routes, together with the lack of obvious kinetic separation among them, further complicates mechanistic interpretation and makes Trp-cage a stringent test for any framework aimed at resolving pathway-specific kinetics.

**FIG. 1.**
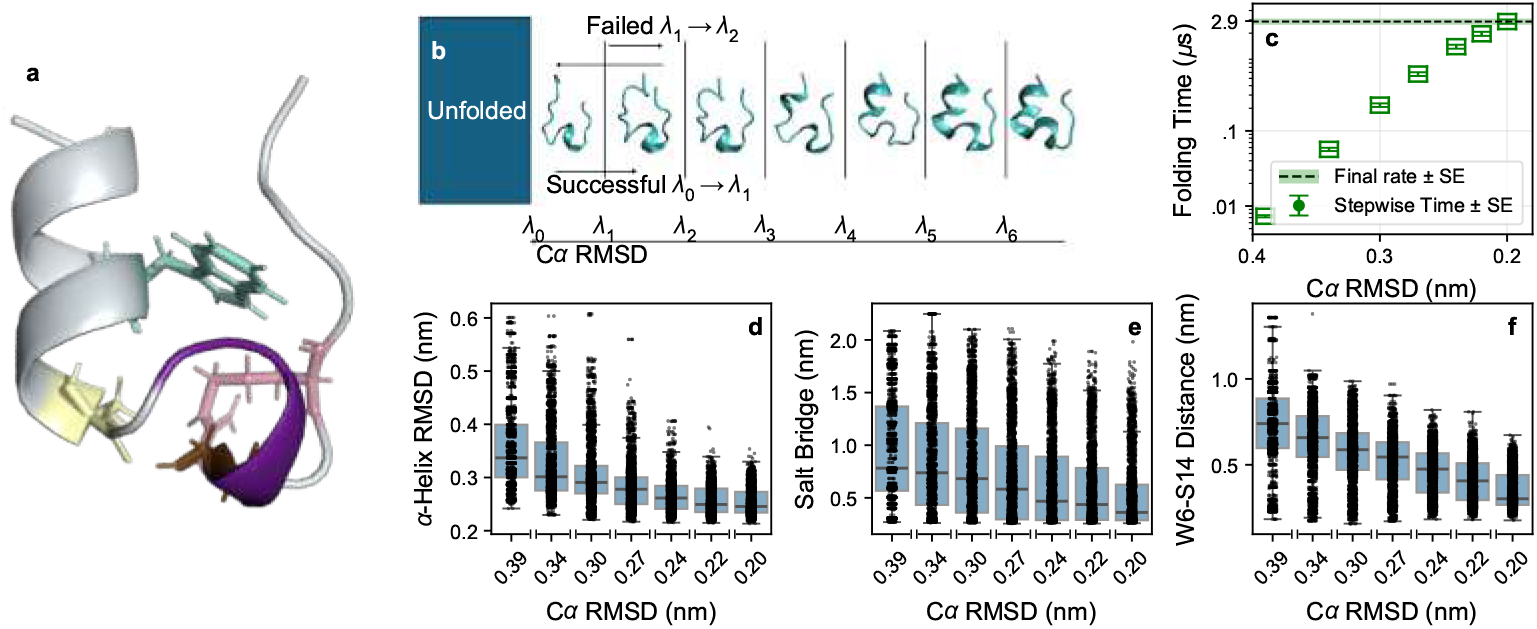
Characterization of the folding kinetics and the evolution of structural order parameters during the folding process. **(a)** PDB structure of the TC5b variant of Trp-cage, with an *α*-helix (grey) and a 3_10_-helix (purple). W6, D9, S14 and R16 residues are depicted in cyan, light yellow, brown, and pink, respectively. **(b)** Schematic representation of the FFS algorithm, involving the initiation of unbiased trajectories within the unfolded basin, and their progression through successive milestones (*λ*_0_to *λ*_6_) toward the folded state via momentum randomization at each milestone. **(c)** Mean waiting times to reach each FFS milestone (depicted by green symbols), with error bars representing the standard errors. The dashed lines and shaded area depict the overall folding time and the standard error, respectively. **(d–f)** Swarm and box plots of **(d)** *α*-helix RMSD relative to the native structure, and **(e–f)** minimum distance between **(e)** D9 and R16 and **(f)** W6 and S14 residues, corresponding to native salt bridge formation and hydrophobic collapse, respectively, for all successful crossings at each milestone.

Here, we introduce a general framework that combines jumpy forward-flux sampling (jFFS) [51] with trajectory-level unsupervised learning [52, 53] to directly compute pathway-resolved kinetics from atomistic simulations. By leveraging ensembles of short, unbiased trajectories, the approach yields thermally averaged rates without relying on biasing potentials, Markovian assumptions, or *a priori* state discretization, and enables a direct decomposition of folding flux into pathway-specific rate contributions with well-defined statistical weights. Applied to the TC5b variant of Trp-cage, we obtain a folding time in near-quantitative agreement with experiments and, from 2,637 statistically representative folding events, identify four dominant pathways distinguished by the temporal ordering of hydrophobic collapse, *α*-helix formation, and salt-bridge stabilization. The resulting mechanistic picture reconciles disparate collapse-first and helix-first scenarios, reveals kinetic degeneracy among some structurally distinct routes, and uncovers a previously unidentified slower frustrated pathway arising from premature salt-bridge formation. More generally, this approach addresses a central limitation in rare-event studies: the inability to convert heterogeneous transition-path ensembles into quantitative, pathway-specific fluxes.

## II. RESULTS AND DISCUSSION

### A. Overall folding kinetics and global trends

To establish the quantitative accuracy of this framework prior to pathway decomposition, we first compute the total folding rate of Trp-cage using jFFS with the *α*-carbon RMSD (*λ*) as the order parameter, which fully distinguishes unfolded and folded states [21]. jFFS partitions folding using a set of C*α* RMSD milestones and reconstructs the rate from the flux out of the unfolded basin and the probability of progressing between successive milestones (Fig. 1b). Because propagation is unbiased, the rate can be estimated from ensembles of short trajectories rather than long continuous simulations, and is generally robust to the choice of order parameter provided it unambiguously separates reactant and product basins, as well as to the placement of intermediate milestones [54].

Applied to the TC5b variant of Trp-cage, this procedure yields a converged folding time of *τ*_fold_ *≈*2.9 *µ*s at 300 K (95% CI: 2.5− 3.2 *µ*s), in near-quantitative agreement with the experimental value of 3 *µ*s at the same temperature [5] (Fig. 1c). This folding time corresponds to the mean first-passage time from the unfolded to folded basin, consistent with two-state kinetic models used in experiments where the unfolded fraction time series is fitted to a single exponential to extract folding and unfolding rates.

To place our results in the context of earlier simulations of Trp-cage folding, we first compare our folding time with the 14 *µ*s value reported in the long-timescale simulations of Lindorff-Larsen *et al*. [15]. The difference should be interpreted in light of two important distinctions between the studies. First, Lindorff-Larsen *et al*. studied a different Trp-cage variant, the 2JOF construct [36], whereas the present work focuses on TC5b. Second, their folding time was estimated from repeated folding and unfolding events as the average residence time in the unfolded state before refolding. This definition measures the refolding latency following an unfolding event, whereas the path-sampling calculation used here yields a thermally averaged mean first-passage time from the unfolded ensemble to the folded basin. Our estimate is also somewhat longer than the folding times reported by Juraszek and Bolhuis [42], who studied the same TC5b variant using transition interface sampling and obtained values of 0.4–1.8 *µ*s, depending on the kinetic definition. These differences are not unexpected, given the known sensitivity of folding kinetics to force fields, sampling protocols, and the specific order parameter used to define interfaces. Diffusion-map analysis has also been used to extract low-dimensional folding coordinates for Trp-cage and to estimate a characteristic folding time by monitoring first arrival into the native basin, yielding an overall ensemble-averaged folding time of 3.73 *µ*s [16]. Importantly, the folding time obtained here is closer to the experimental temperature-jump estimate for TC5b than in all such earlier studies, providing an independent benchmark for the present calculations.

To elucidate the folding mechanism, we examine the evolution of key structural order parameters averaged over crossings at successive milestones. The *α*-helix RMSD decreases steadily (Fig. 1d), hydrophobic packing– quantified by the Trp6–Ser14 (W6–S14) distance– tightens progressively (Fig. 1f), and the Asp9– Arg16 (D9–R16) salt bridge forms predominantly at later stages (Fig. 1e). Global descriptors, including solvent-accessible surface area (Fig. S1a) and radius of gyration (Fig. S1b), show similarly continuous declines. However, these smooth averages mask broad– and in some cases bimodal– distributions at early milestones (Fig. S2), indicating that *α*-helix formation, hydrophobic collapse, and electrostatic stabilization occur on overlapping timescales, and none uniquely commits the system to the folded state. Instead, folding proceeds through distinct sequences of partially coupled rearrangements, with multiple pathways emerging early and coexisting throughout the transition region.

### B. Identifying physically meaningful pathways

To resolve mechanistic heterogeneity in the folding ensemble, we analyze FFS folding events using dimensionality reduction. Each folding event ℱ_*k*_ = {𝒞_*k*,0_, 𝒞_*k*,1_, *…*, 𝒞_*k*,6_} is a milestone-discretized trajectory, where 𝒞_*k,j*_ is the endpoint of an unbiased trajectory initiated from 𝒞_*k,j−*1_. Collectively, these folding events encode the temporal ordering of structural rearrangements, but their high dimensionality and intrinsic correlations make it infeasible to directly group them into distinct interpretable pathways through visual inspection or simple structural analysis. Standard dimensionality-reduction approaches, including principal component analysis and a conventional autoencoder [55], capture local variance but do not yield clear mechanistic separation (Fig. S3), motivating a representation that preserves the dynamical evolution of entire trajectories.

We therefore train a variational autoencoder [52, 53] (VAE) on *α*-carbon positions across all folding events to learn a low-dimensional representation of entire folding trajectories (Fig. 2a). Embedding trajectories, instead of configurations, proves essential as it preserves ancestral relationships and captures correlated structural evolution, thereby enabling separation of pathways based on their dynamical progression rather than static structural similarity. Here, the embedding is used to classify complete FFS transition events, in contrast to earlier applications of deep learning that identify molecular representations or kinetic coordinates for constructing biased collective variables [56, 57] or Markovian kinetic models [58, 59]. The probabilistic bottleneck promotes a smooth non-degenerate latent space that maintains pathway diversity while avoiding artificial clustering. The resulting two-dimensional latent space (Figs. 2b–d) forms well-separated clusters that correlate with early-milestone order parameters– *α*-helix RMSD (Fig. 2b),W6–S14 distance (Fig. 2c), and the D9–R16 salt bridge distance (Fig. 2d)– without explicit supervision, indicating that the model autonomously identifies the structural features most relevant to folding (see Figs. S4 and S5 for training protocol and order parameter overlays across all milestones, respectively).

**FIG. 2.**
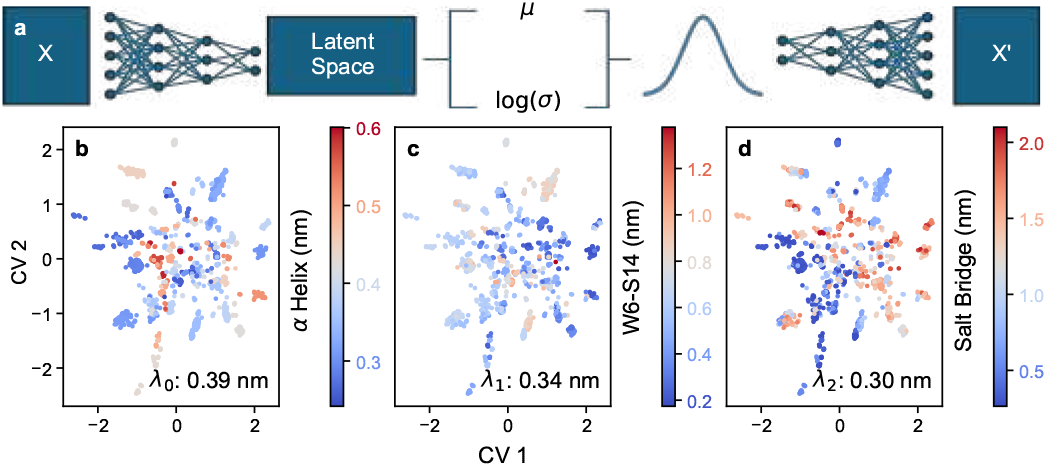
Variational autoencoder architecture, and latent space visualization based on physical order parameters. **(a)** Diagram illustrating the VAE architecture, with a loss function comprised of mean squared error (MSE) and Kullback–Leibler (KL) divergence terms. **(b–d)** The obtained latent space colored in terms of physical order parameters at successive milestones: **(b)** *α*-helix RMSD at *λ*_0_= 0.39 nm, **(c)** W6–S14 distance at *λ*_1_= 0.34 nm, and **(d)** D9–R16 salt bridge distance at *λ*_2_= 0.30 nm

To obtain physically interpretable folding pathways, we apply density-based clustering [60] in the latent space, yielding ∼30 micro-clusters (Fig. S6). Four dominant clusters account for nearly half of all folding events. The remaining low-population clusters are therefore assigned to the nearest major clusters using *α*-helix RMSD, W6–S14 distance, and D9–R16 salt bridge values at *λ*_0_ (Fig. 3f, Section S2). Thus, the VAE reveals the trajectory-level organization of the transition ensemble, whereas the final pathway definitions are expressed in terms of physically interpretable *λ*_0_ descriptors so that pathway-specific fluxes can be computed reproducibly. The coarse-grained pathways, whose populations and milestone evolution are summarized in Figs. 3b and 4a, respectively, reveal distinct sequences of structural rearrangements defined using descriptors not employed during training. Extensive validation–including held-out retraining, latent-space scans, and clustering stability analyses–confirms the robustness of this classification (Section S2, Figs. S4, S6).

**FIG. 3.**
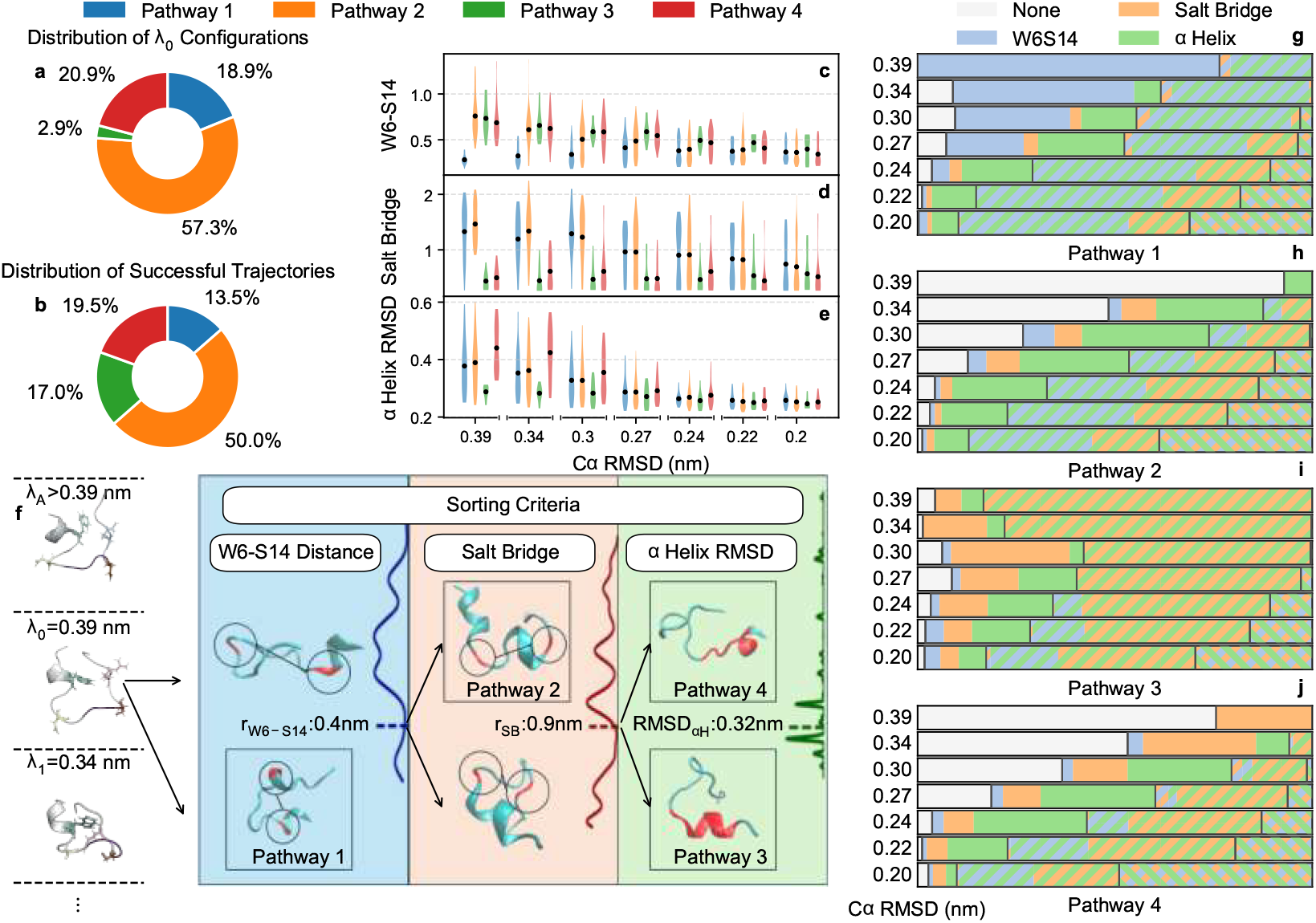
Pathway decomposition into structurally and kinetically distinct folding routes. **(a–b)** Distribution of configurations assigned to each pathway at **(a)** *λ*_0_= 0.39 nm and **(b)** *λ*_6_= 0.20 nm. **(c–e)** Pathways differ in key structural descriptors, namely **(c)** W6–S14 hydrophobic contact distance, **(d)** D9–R16 salt bridge distance, and **(e)** *α*-helix RMSD. **(f)** Diagram illustrating the process to sort configurations at *λ*_0_ into distinct pathways, with embedded histograms indicating cutoffs in each property used for clustering. (**g-j**) Prevalence of secondary and tertiary structural elements at each milestone within each pathway. We use an *α*-helix RMSD cutoff of 0.29 nm to detect an *α*-helix, and W6–S14 and D9–R16 cutoffs of 0.4 and 0.46 nm to identify W6–S14 contacts and salt bridges, respectively. These cutoffs differ from those utilized for clustering in (**f**). Hashed portions correspond to simultaneous existence of multiple structural elements.

### C. Pathway-specific folding times

We next perform the central step of the framework: direct decomposition of the total folding flux into pathway-specific rate contributions. The pathway classification from Fig. 3f is applied to all configurations at the first milestone, *λ*_0_, regardless of whether they reach the folded basin, with the resulting entry fractions shown in Fig. 3a. These fractions quantify how often the unfolded ensemble enters each pathway; the corresponding flux contributions also depend on the conditional probability of reaching the folded basin after pathway entry. For each pathway *p*, the rate is computed as *R*_*p*_ = Φ_0_ Π*j P* (*λ*_*j*+1_|*λ*_*j*_; *p*), where *P* (*λ*_*j*+1_ *λ*_*j*_; *p*) is the conditional probability for configurations at *λ*_*j*_ whose ancestors at *λ*_0_ belong to *p* (see SI Section S3). The folding time, *τ*_*p*_ = 1*/R*_*p*_, represents the mean first-passage time *conditioned on entry* into pathway *p*, disentangling intrinsic kinetics from the pathway’s selection probability in the unfolded ensemble. In this representation, pathways are treated as ancestry-conditioned kinetic sub-ensembles (defined per entry criteria of Section II B) rather than absorbing dynamic states. Unlike prior approaches that identify pathways without quantifying their kinetic contributions, this decomposition assigns a rate to each pathway consistent with the total folding flux.

The pathway-specific folding times span over an order of magnitude (Figs. 5a–b). Each rate is accompanied by a conditional survival probability profile (Fig. 5c), defined as the cumulative product of transition probabilities across remaining milestones. As established previously [54], the survival probability provides an FFS-based proxy for commitment by quantifying the likelihood that trajectories initiating from a given milestone will reach the folded state. These profiles therefore indicate when trajectories within each pathway become kinetically committed and provide a natural ordering of structural events along decreasing C*α* RMSD.

### D. Pathway-specific mechanisms

Having established a physically interpretable partitioning of the folding ensemble and quantified pathway-specific rates, we next examine the sequence of structural events in each pathway (Figs. 3g–j) to identify mechanistic signatures underlying their distinct kinetics.

#### Pathway 1: Early non-native collapse

This pathway (*τ*_fold_ = 3.5 *±*0.5 *µ*s) features early compaction, indicated by small W6–S14 distances at *λ≈* 0.39–0.34 nm (Fig. 3c) that persists at later milestones (Fig. 3g), also evident in a representative folding event shown in Fig. 4b. Despite this collapse, the conditional survival probability remains low until *λ≈* 0.24 nm, indicating delayed commitment, which occurs only after *α*-helix formation and D9–R16 salt bridge establishment (Figs. 3d–e,g; Fig. 4b; SI Movie S1). These observations reveal a decoupling between early compaction and native topology acquisition: folding requires escape from a compact, non-native intermediate followed by contact reorganization.

**FIG. 4.**
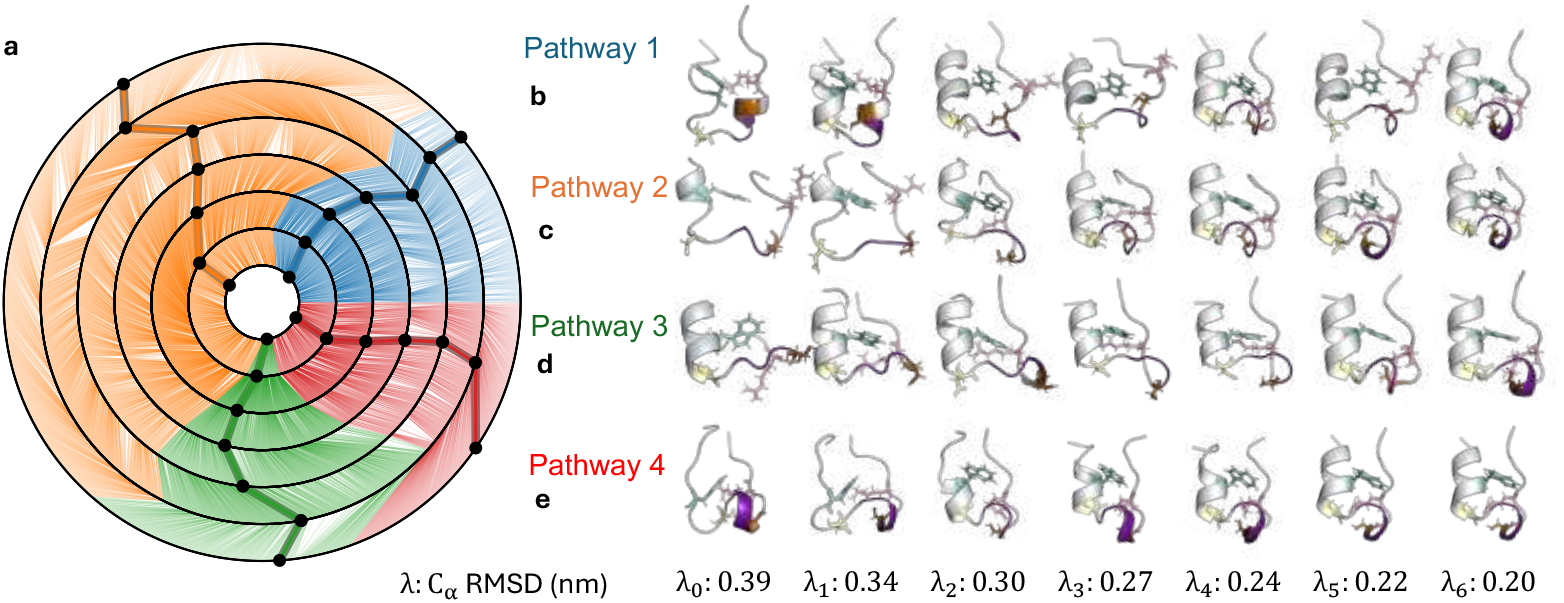
Representative folding events. (**a**) Diagrammatic representation of all reactive folding events. Each node represents a unique configuration, with connecting lines corresponding to successful FFS trajectories. Each concentric circle corresponds to an FFS milestone, with innermost circles corresponding to earlier milestones. (**b–e**) Instantaneous snapshots of representative folding events in each pathway, with *α*-helix (gray), 3_10_-helix (purple), W6 (cyan), D9 (yellow), S14 (brown) and R16 (pink).

This collapse-first mechanism is consistent with prior atomistic [41] and coarse-grained [40] studies reporting early hydrophobic collapse. However, its delayed commitment to the folded basin suggests that folding barriers arise from late-stage topological alignment rather than initial collapse. Notably, this pathway accounts for only 13.5% of folding events, whereas earlier transition-path-sampling work identified it as the dominant route, comprising ∼80% of sampled trajectories [41]. The difference likely reflects the enhanced statistical breadth of the present transition ensemble and the use of trajectory-level classification to assign pathway weights.

#### Pathway 2: Helix-first pathway

Accounting for roughly half of all folding events, this pathway folds in *τ*_fold_ = 3.1 ± 0.3 *µ*s. Parent configurations at *λ*_0_ lack substantial secondary or tertiary structure (Fig. 3h), yet *α*helix formation occurs early, followed by later hydrophobic collapse and salt-bridge formation (Figs. 3c-e; Fig. 4c; SI Movie S2). This mechanism is consistent with prior studies emphasizing the dominance of early helix formation [38, 44, 45].

Despite distinct structural sequences, this pathway exhibits folding times comparable to the collapse-first route, illustrating *kinetic degeneracy*; different molecular rearrangements yield indistinguishable kinetics. In particular, neither early helix formation nor hydrophobic collapse alone confers a kinetic advantage for folding. Such degeneracy may enhance evolutionary robustness by distributing folding flux across multiple routes, reducing sensitivity to mutations or external perturbations.

#### Pathway 3: Early helix-salt bridge pathway

This is the fastest pathway (*τ*_fold_ = 0.41 ± 0.08 *µ*s), characterized by early *α*-helix formation and salt-bridge stabilization, followed by hydrophobic collapse (Figs. 3c–e; Fig. 4d; SI Movie S3). Despite its speed, it comprises only ∼17% of folding events, likely because accessing this route carries a substantial entropic cost: in the unfolded ensemble, only 2.9% of *λ*_0_ configurations simultaneously exhibit both the helical and salt-bridge features required to initiate it (Figs. 3a,b). Although this pathway is initiated by a small fraction of *λ*_0_ configurations, its enhanced transition probabilities persist across subsequent milestones (Table S2), yielding a conditional folding time that remains well separated from the other pathways within uncertainty. Consequently, once engaged, this pathway has a higher survival probability, causing it to make up an increasing fraction of trajectories at later folding milestones (Fig. 5c).

**FIG. 5.**
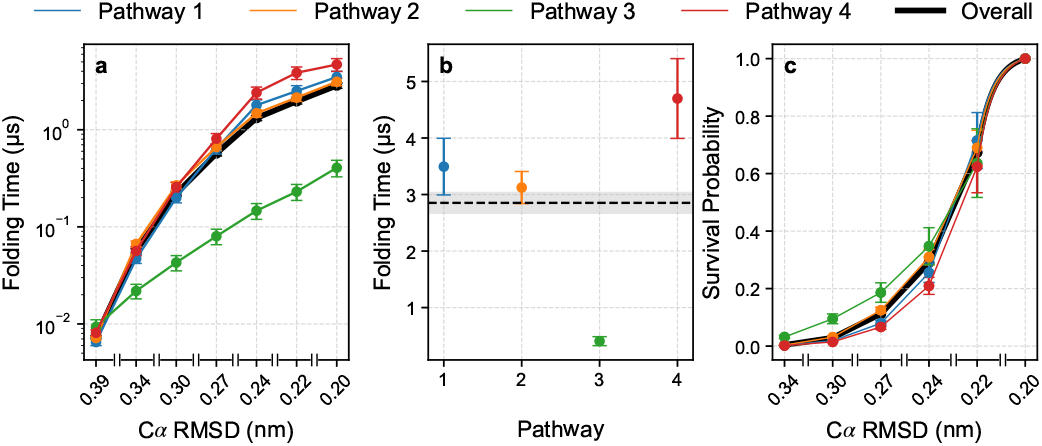
Pathway-specific folding times. (**a**) Cumulative total (black) and pathway-specific mean first-passage time vs. C*α* RMSD, with (**b**) overall folding times revealing marked differences among pathways. The grey shade in (**b**) corresponds to the mean and the standard error in total folding time. (**c**) Total (black) and pathway-specific (colored) survival probability vs. C*α* RMSD.

While superficially similar to the helix-first pathway, its faster kinetics show that helix formation alone offers limited kinetic advantage without concurrent electrostatic stabilization, and reveal substantial kinetic heterogeneity within previously classified helix-first pathways [38, 41, 44, 45].

#### Pathway 4: Kinetically frustrated pathway with off-pathway detours

This is the slowest route, *τ*_fold_ = 4.7*±* 0.7 *µ*s, with consistently lower survival probabilities until late milestones (Fig. 5c). It exhibits delayed hydrophobic collapse, persistently high *α*-helix RMSDs (until *λ* ≥ 0.27 nm), and non-monotonic salt-bridge distances reflecting early formation, rupture, and reformation (Figs. 3c-e,j; Fig. 4e; SI Movie S4). Together, these features indicate prolonged residence in weakly committing, “misfolded” states, where premature salt-bridge formation delays productive folding by diverting trajectories into off-pathway intermediates requiring extensive reorganization before native topology is adopted (Fig. 4e). Such frustrated routes are broadly relevant to protein misfolding and aggregation disorders.

Unlike the fastest pathway, configurations are stabilized in turn-rich, non-helical geometries at *λ*_0_ despite a pre-formed salt bridge, consistent with experimental observations of salt bridge-stabilized intermediates [37]. Productive folding therefore requires electrostatic reconfiguration: the premature salt bridge must dissociate to permit helix nucleation and core repacking, and then reform in the native register to stabilize the folded structure. The need to break and re-form a strong interaction explains the order-of-magnitude slowdown (in comparison to Pathway 3) and highlights the sensitivity of folding kinetics to subtle early structural differences.

Although this pathway has not been previously identified explicitly, prior studies support off-pathway frustration in Trp-cage folding, including mispacked intermediates [43], frustrated detours [41], and a metastable compact basin with two partially-formed hydrophobic sub-cores linked by a mis-formed salt bridge [17]. The delayed rise in survival probability observed here is a kinetic signature of such frustration, consistent with kinetic partitioning [4, 61–64], in which trajectories split into fast-committing and slower subpopulations trapped in metastable intermediates.

### E. Broad kinetic and mechanistic observations

The pathway-resolved decomposition reveals several general features of folding. Approximately 50% of events proceed via the helix-first pathway (Fig. 3b), with the remaining flux distributed among the other routes. The helix–salt bridge pathway, though initially underrepresented, becomes progressively enriched as *λ* decreases, reaching ∼17% at the final milestone, whereas the other pathways are progressively depleted. These observations demonstrate that structural prevalence is a poor proxy for kinetic potency– a distinction accessible only through explicit path sampling.

A second insight is the central role of *α*-helix formation across all pathways. Even in collapseor salt bridge-dominated routes, *α*-helices emerge early (Figs. 3h,j), and concurrent formation of the W6–S14 contact and D9–R16 salt bridge is rare without a pre-formed *α*-helix (Figs. 3h-j). Tertiary contacts alone are therefore insufficient to stabilize the folded state. Together with the comparable folding times of the collapse-first and helix-first routes, this observation suggests that kinetic advantage arises from coordinated ordering of secondary and tertiary elements rather than from any single early structural event.

The qualitative features of the four pathways, each reflecting a distinct ordering of key secondary-structure elements and tertiary contacts in Trp-cage, raise an interesting possibility: at a coarse-grained level, plausible folding routes can be viewed as different orderings of a limited set of structural motifs. Because the number of such permutations grows exponentially with the number of underlying motifs, this view suggests a potential combinatorial proliferation of candidate pathways for larger proteins, reminiscent of *Levinthal’s paradox* [62]. However, the observation that only a small subset of these plausible routes is kinetically selected–four in the present work–is consistent with the folding-funnel picture, in which the energy landscape channels conformational search toward a limited number of productive pathways.

As noted above, the pathway-specific folding times are conditional on λ_0_ configurations being pre-assigned to a given pathway and do not reflect the probability of accessing that pathway. The physically relevant folding flux through pathway *p* is thus given by *f*_*p*_/*τ*_*p*_, where *f*_*p*_ is the fraction of λ_0_ configurations pre-assigned to pathway *p*. With this correction, the sum of pathway fluxes, (3.5 ± 0.2) × 10^5^ s^−1^, matches the total folding flux, 1/*τ*_fold_ = (3.5 ± 0.2) × 10^5^ s^−1^ (Table S1).

As discussed in Section II C, ancestry-conditioned subensembles are not identical to pathways identified from the completed transition ensemble (Fig. 3b). The two representations become equivalent when the pathway label assigned at λ_0_ captures the kinetic identity of the completed folding event, so that trajectories within a given ancestry-defined sub-ensemble do not subsequently partition into distinct routes with different folding probabilities. This equivalence can be assessed by comparing the flux fractions inferred from ancestry-conditioned folding times (Table S1) with those obtained by assigning completed folding events to the four clustered pathways (Fig. 3b). In the limit of adequate sampling and kinetically coherent pathway labels, these two estimates should converge to the same pathway flux fractions. Their agreement within statistical uncertainty for Pathways 1–3 supports the ancestry-conditioned decomposition. Pathway 4 is the only notable deviation, consistent with its non-uniform survival profile, in which weak early commitment is followed by enhanced late-stage transition probabilities.

Unlike prior approaches relying on biasing potentials or long unbiased simulations, our framework assigns statistically robust kinetic weights to each pathway, yielding pathway-resolved rates and fluxes with rigorous uncertainties. By identifying four physically interpretable routes and quantifying their contributions, we resolve longstanding ambiguities in Trp-cage folding and reveal a kinetically frustrated pathway that had not previously been identified directly.

## III. CONCLUSIONS AND FUTURE OUTLOOK

Here, we introduce a framework that combines forward-flux sampling with trajectory-level embedding to estimate pathway-specific folding times from unbiased atomistic simulations. Unlike prior enhanced-sampling and long-trajectory studies [15, 16, 41, 48], which identified plausible folding sequences without quantifying their frequencies or kinetics, our approach assigns thermally averaged statistical weights to physically interpretable pathways, directly linking structural evolution to kinetic relevance, and enabling quantitative comparison of competing mechanisms.

The framework is both robust and general. Because FFS computes rates directly from initial fluxes and transition probabilities, it eliminates the need for reweighting, bias removal, *a priori* state discretization, or Markovian assumptions. Its tolerance to suboptimal order parameters [54], combined with efficient parallelization and compatibility with driven dynamics [65, 66], makes it well suited for a wide range of biomolecular rare events, including nonequilibrium processes such as field- and strain-induced conformational changes [67–69] and driven solute transport through biological channels [70–72].

The approach should extend beyond miniproteins to folding systems with multiple nucleation sites [73], fold-switching proteins [74], and repeat proteins with multiple initiation points [75]. For larger proteins, however, hierarchical folding, long-lived intermediates, and increased memory effects may require multiple rate calculations and more careful sampling of the unfolded basin. The practical limiting step is therefore not the formal rate decomposition, but the construction of an order parameter and basin ensemble that capture all kinetically relevant entry routes. Even in such cases, the framework remains applicable provided the starting ensemble is adequately sampled.

By resolving total folding flux into structurally interpretable pathways, our approach provides a roadmap for dissecting how sequence, force field, and environmental factors—temperature, ionic strength, small molecules, and crowding—reshape folding kinetics, particularly when misfolding competes with productive folding. Identifying distinct misfolding routes and their structural checkpoints further reveals how such perturbations bias early commitment toward deleterious pathways, while highlighting interactions that act as kinetic safeguards against mispacked collapse.

The same strategy extends naturally to ligand binding, conformational switching, and allostery, where competing pathways govern function [76–78]. Embedding trajectories from different variants and conditions into a shared latent space enables direct comparison of mechanisms, identification of bottlenecks and metastable intermediates, and closer integration with temperature-jump and single-molecule experiments.

In summary, converting transition-path ensembles into pathway-resolved rate contributions provides the missing kinetic context for systems with multiple competing mechanisms, and offers a route to understanding and ultimately controlling kinetic competition in complex chemical and biomolecular systems.

## MATERIALS AND METHODS

### Molecular Dynamics Simulations

Protein and water molecules are modeled using the Amber03w [79] and TIP4P/2005 [80] force-fields, respectively. All MD simulations are performed in the isothermal–isobaric (NPT) ensemble using GROMACS [81]. The equations of motion are integrated with the leap-frog algorithm with a 2-fs time step. Temperature and pressure are controlled using the Nosé -Hoover [82, 83] thermostat (0.2ps time constant) and the Parrinello–Rahman [84] barostat (2-ps time constant). Long-range electrostatics are treated with the smooth particle-mesh Ewald [85] (PME) method, employing a 1-nm cutoff for short-range interactions, and long-range dispersion corrections are applied to both potential energy and pressure. The geometries of water molecules and rigid covalent bonds in Trp-cage are constrained using Settle [86] and LINCS [87], respectively. The cubic simulation box contains 2,910 water molecules, a single Trp-cage chain, and a chloride ion to neutralize the Trp-cage’s net charge at pH 7.

### Folding Rate Calculations

Folding rates are calculated using jFFS [51] with Cα RMSD to the native structure [35] (PDB ID 1L2Y) as the order parameter. Configuration space is partitioned using level sets of the order parameter– called milestones– and the folding rate is estimated iteratively by multiplying Φ_0_, the flux of trajectories initiated in the unfolded state and reaching the first milestone, λ_0_, and *P*(λ_*j*+1_| λ_*j*_)’s, the probabilities of transitioning from each milestone to the subsequent one (Fig. 1b). The initial flux is estimated from short unbiased trajectories that repeatedly recross λ_0_, while transition probabilities are evaluated by launching ensembles of momenta-randomized, unbiased trajectories from milestone-crossing configurations at the preceding iteration. All trial trajectories are conducted at 300 K and 1 bar, with the order parameter monitored every 1 ps. The starting configurations for basin exploration simulations are obtained by solvating and equilibrating 250 representative unfolded structures from a prior [21] REMD simulation (72 replicas, from 210 K to 496.5 K at 1 bar). Basin exploration is conducted with λ_basin_ = 0.55 nm and λ_0_ = 0.39 nm, leading to the collection of 980 configurations at λ_0_ over 7.16 *µ*s, corresponding to 1.08 *×*10^4^*τ* where *τ* is the order parameter autocorrelation timescale (See Section S4 and Fig. S7 for further details). Six FFS iterations are conducted afterwards, with the target milestone of each iteration set based on the OP distribution of the prior iteration, following the method of Ref. 51. A minimum of 2,500 crossings are obtained per iteration.

Error bars in transition probabilities are computed using the landscape variance approach of Allen *et al*. [88]. Total and pathway-specific folding times, as well as associated transition probabilities are summarized in Tables S1 and S2, respectively.

### Transition Ensemble Analysis using Variational Autoencoders

VAE [53] is trained on C*α* positional features from 2,637 folding events obtained in FFS. The encoder ingests tensors of C*α* coordinates across seven FFS milestones, resulting in a tensor shape of (2637, 7, 20, 3), and produces a two-dimensional latent embedding (2637, 2). Unlike standard autoencoders, the VAE includes a probabilistic layer to foster a continuous latent space. The loss function comprises both a mean-squared error (MSE) term and a Kullback–Leibler (KL) divergence term, with the KL term regularizing the latent distribution. To balance the opposing effects of MSE and KL losses, a cyclical KL-weight schedule is employed, ramping up and resetting over ten cycles of 500 epochs each. This approach yields a compact yet expressive latent space (see Figs. 2b-d), which we partition into approximately 30 micro-clusters using density-based clustering. However, four principal clusters account for about half the folding events, and all remaining minor clusters are assigned to the four major ones discussed in the text. A detailed discussion and justification of the employed classification approach is given in Section S2. To avoid overfitting, training was repeated with an 80/20 training/test split of the folding events, yielding statistically indistinguishable latent spaces and losses.

## DATA AVAILABILITY

All data needed to evaluate the conclusions in the paper are present in the paper and/or the Supporting Information. Simulation input files, processed data, representative configurations, and analysis scripts will be deposited in a public repository before publication. Full raw trajectory files will be available from the corresponding author upon reasonable request because of their large size.

## ACKNOWLEDGMENTS

This work was supported by the Alfred P. Sloan Foundation. B.U. acknowledges the Anderson Postdoctoral Fellowship from Yale University and Bogaziçi University Research Fund (BAP Project No. 25A05P5). M.A.L. acknowledges support from the Program in Physical and Engineering Biology (PEB) at Yale University. These calculations were performed at the Yale Center for Research Computing. This work used Stampede through allocation CHM240063 from the Advanced Cyberinfrastructure Coordination Ecosystem: Services & Support (ACCESS) program, which is supported by NSF Grants #2138259, #2138286, #2138307, #2137603, and #2138296.

