## Supporting Information for "Pathway-resolved flux decomposition reveals hidden kinetic hierarchy in protein folding"

(Dated: May 20, 2026)

### S1. ALTERNATIVE DIMENSIONALITY REDUCTION APPROACHES

Prior to utilizing the variational autoencoder (VAE), we explore two alternative dimensionality reduction strategies. First, we apply conventional principal component analysis (PCA). The input vectors are constructed from five structural order parameters–$R_g$, SASA, $\alpha$-helix RMSD, the W6–S14 distance, and the D9–R16 salt bridge distance– evaluated along individual folding events. This representation maps the original $(N, 5, 7)$ dataset onto a two-dimensional $(N, 2)$ embedding, where $N = 2{,}637$ denotes the number of folding events.

As shown in Figs. S3a-c, the PCA projection fails to produce a clear separation of folding pathways and does not resolve distinct structural features. This outcome reflects the intrinsic linearity of PCA, which limits its ability to capture the nonlinear correlations that govern the underlying folding mechanisms.

As a second approach, we implement a conventional autoencoder [55] trained jointly on the $\alpha$-carbon Cartesian coordinates (as in the VAE) and the selected structural order parameters outlined above. The positional data are first processed through three one-dimensional convolutional layers with strides of 8, 16, and 32. The resulting feature maps are flattened and passed through two fully connected layers, reducing the input from $(N, 7, 20, 3)$ to a latent representation of size $(N, 16)$. In parallel, the order parameters are encoded using three fully connected layers designed to yield an output of the same dimensionality. The two encoded representations are concatenated and supplied to a third encoder of similar architecture, producing a shared latent space.

Decoding proceeds along two separate branches to reconstruct the original inputs. For the positional data, two large fully connected layers are followed by two 2D convolutional layers to recover the original tensor shape. The order parameters are reconstructed using an analogous decoder, with modified convolutional specifications to match their input dimensions.

Despite its architectural flexibility, this autoencoder does not yield a clear separation between folding pathways, nor did it reveal a meaningful spatial organization of the order parameters in latent space (Figs. S3d-f). Furthermore, the requirement that the network be trained explicitly on predefined order parameters is suboptimal, as it introduces prior bias into a procedure intended to uncover intrinsic structural organization.

### S2. ASSIGNMENT OF MINOR CLUSTERS TO THE FOUR MAJOR CLUSTERS

We employ density-based clustering to partition the full folding ensemble into distinct mechanistic clusters. Four major clusters emerge (Fig. S6a), collectively accounting for more than 40% of all folding trajectories (Fig. S6d). These dominant pathways are compared using the statistical distributions of the W6–S14 distance (Fig. S6i), the D9–R16 salt bridge distance (Fig. S6h), and the $\alpha$-helix RMSD (Fig. S6g) evaluated at $\lambda_0$, which collectively yield the classification criteria summarized in Fig. 3f.

To assign minor clusters onto the four major pathways, we consider two assignment strategies. In the first approach (Fig. S6b,e,j–l), mean values of the relevant order parameters are computed for each minor cluster, which is subsequently assigned to the major pathway whose classification criteria are satisfied by these averages. This procedure yields pathways that remain geometrically contiguous in the latent space (Fig. S6b), although a small number of individual folding events display order parameter values inconsistent with the assigned classification.

In the second approach (Fig. S6c,f,m–o), assignments are performed at the folding event level using instantaneous order parameter values. This approach introduces a limited number of latent-space outliers (Fig. S6c)– folding events classified within a pathway but embedded among those of another– yet such cases constitute only 198 folding events (accounting for only 7.5% of the reactive trajectory ensemble) and produce no discernible changes in the resulting distributions (Figs. S6e–f) and violin plots (Figs. S6j–l vs. S6m–o). We therefore adopt the trajectory-level assignment strategy, as it enables the formulation of well-defined entry criteria suitable for rigorous computation of pathway-specific folding rates.

*

### S3. PATHWAY-SPECIFIC FOLDING RATES

To compute the folding time associated with each pathway, we partition the configurations at $\lambda_0$ into four clusters according to the criteria described in Fig. 3f, effectively decomposing the ensemble of first-milestone crossings into four non-overlapping regions. (This is akin to treating each pathway as an ancestry-conditioned subensemble, and not an absorbing dynamic state.) A partial flux associated with pathway $p$ can then be defined as

$$R_p = \frac{N_{0,p}}{t_{\text{basin}}} \prod_{k=0}^{N-1} P(\lambda_{k+1}|\lambda_k; p), \tag{S1}$$

where $t_{\text{basin}}$ denotes the cumulative duration of unbiased MD trajectories conducted within the unfolded basin, and $N_{0,p}$ is the number of crossings at $\lambda_0$ assigned to pathway $p$. By construction, $N_0 = \sum_p N_{0,p}$, with $N_0$ the total number of crossings at $\lambda_0$. The quantities $P(\lambda_{k+1}|\lambda_k; p)$ represent the conditional transition probabilities for configurations at interface $\lambda_k$ that originate from a $\lambda_0$ configuration assigned to pathway $p$.

With this formalism, $\sum_p R_p \to R$ in the limit of infinite sampling ($t_{\text{basin}} \to \infty$), such that each $R_p$ encodes both the probability of entering pathway $p$ at $\lambda_0$ and the subsequent folding rate along that pathway. In the main text, however, we report the mean first passage time (MFPT) to the folded state *conditioned* on entry into a given pathway. Accordingly, (S1) must be rescaled by the probability of entering pathway $p$ at $\lambda_0$, namely $N_{0,p}/N_0$, yielding

$$\begin{aligned} R_{p,\text{cond}} &= \frac{R_p}{N_{0,p}/N_0} = \frac{N_0}{t_{\text{basin}}} \prod_{k=0}^{N-1} P(\lambda_{k+1}|\lambda_k; p) \\ &= \Phi_0 \prod_{k=0}^{N-1} P(\lambda_{k+1}|\lambda_k; p) \end{aligned} \tag{S2}$$

where $\Phi_0 = N_0/t_{\text{basin}}$ is the total flux through $\lambda_0$. The pathway-specific MFPT then follows directly as $\tau_p = 1/R_{p,\text{cond}}$.

### S4. ORDER PARAMETER AUTOCORRELATION FUNCTION

To determine an appropriate level of sampling of the unfolded basin and to select a suitable order parameter sampling window, we compute the autocorrelation function of the order parameter, $C_\lambda(t)$, defined as,

$$C_\lambda(t) = \frac{\langle [\lambda(t) - \overline{\lambda}][\lambda(0) - \overline{\lambda}] \rangle}{\langle [\lambda(0) - \overline{\lambda}]^2 \rangle}.$$

Here, $\overline{\lambda}$ denotes the mean value of the order parameter along a given trajectory. To estimate the uncertainty in the autocorrelation function, $C_\lambda(t)$ is computed separately for each trajectory, and the mean and standard error across trajectories are evaluated for every $t$. The resulting autocorrelation function is shown in Fig. S7.

As a general heuristic, the starting (unfolded) basin should be sampled for at least$\sim 10^4\tau$, where $\tau$ is the autocorrelation time obtained from $C_\lambda(t)$. From the $C_\lambda(t)$ depicted in Fig. S7, we estimate a relaxation time of $\tau = 664$ ps. Our 7.16-$\mu$s exploration of the unfolded basin therefore corresponds to $1.08 \times 10^4\tau$, which is sufficient to generate a diverse ensemble of $\lambda_0$ configurations. Furthermore, the order parameter sampling window of 1 ps used in this work is between two to three orders of magnitude smaller than $\tau$, consistent with recommendations in our earlier work [89].

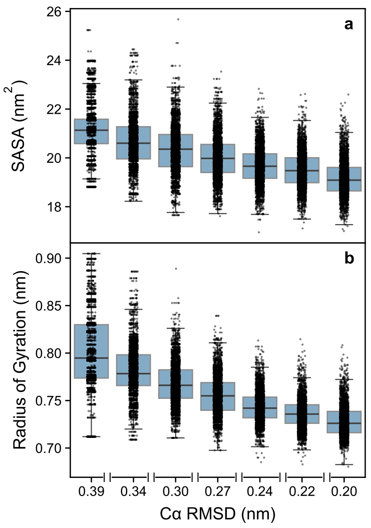


FIG. S1. *Evolution of order parameters not utilized for clustering along the folding pathway.* Swarm plots of (a) solvent accessible surface area and (b) radius of gyration.

TABLE S1. Total and pathway-specific folding times and fluxes, with $\boldsymbol{\tau}_0 = 1/\boldsymbol{\Phi}_0 = 7.30468 \pm 0.24209$ ns. Pathway-specific folding times, $\boldsymbol{\tau}_p$'s, are conditional on entry into pathway $p$ at $\boldsymbol{\lambda}_0$, whereas the physically relevant flux contribution is $f_p R_p = f_p/\boldsymbol{\tau}_p$, with $f_p$ values given in Fig. 3a. Accordingly, $\sum_p f_p R_p = 0.3500 \pm 0.0248\,\mu\text{s}^{-1}$ is statistically indistinguishable from $R_{\text{total}}$.

| Pathway, $p$ | Cumulative probability, $P(\lambda_6\|\lambda_0; p)$ | Folding time, $\tau_f$ [$\mu$s] | $f_p$ | $f_p R_p$ [$\mu\text{s}^{-1}$] | $f_p R_p / \sum_{p'} f_{p'} R_{p'}$ |
|---|---|---|---|---|---|
| total | $0.0025607 \pm 0.0001649$ | $2.8526 \pm 0.1837$ | 1.000 | $0.3506 \pm 0.0226$ | – |
| 1 | $0.0020909 \pm 0.0002996$ | $3.4936 \pm 0.5138$ | 0.189 | $0.0541 \pm 0.0080$ | $0.1543 \pm 0.0248$ |
| 2 | $0.0023388 \pm 0.0002131$ | $3.1233 \pm 0.3027$ | 0.573 | $0.1835 \pm 0.0178$ | $0.5233 \pm 0.0609$ |
| 3 | $0.0179865 \pm 0.0034871$ | $0.4061 \pm 0.0787$ | 0.029 | $0.0714 \pm 0.0138$ | $0.2037 \pm 0.0416$ |
| 4 | $0.0015547 \pm 0.0002336$ | $4.6985 \pm 0.7061$ | 0.209 | $0.0445 \pm 0.0067$ | $0.1269 \pm 0.0207$ |

TABLE S2. Total and pathway-specific transition probabilities. Here, $N_c$, $N_t$ and $N_s$ correspond to the total number of starting configurations, trial trajectories, and successful crossings, respectively.

| Pathway, $p$ | $i$ | $\lambda_i$ [nm] | $\lambda_{i+1}$ [nm] | $N_c$ | $N_t$ | $N_s$ | $P(\lambda_{i+1}\|\lambda_i; p)$ |
|---|---|---|---|---|---|---|---|
| total | 0 | 0.39 | 0.34 | 980 | 23,713 | 3,053 | $0.12875 \pm 0.00645$ |
| total | 1 | 0.34 | 0.30 | 3,053 | 23,905 | 6,157 | $0.25756 \pm 0.00587$ |
| total | 2 | 0.30 | 0.27 | 6,157 | 14,236 | 5,495 | $0.38599 \pm 0.00685$ |
| total | 3 | 0.27 | 0.24 | 5,495 | 10,957 | 4,732 | $0.43187 \pm 0.00789$ |
| total | 4 | 0.24 | 0.22 | 4,732 | 7,627 | 5,137 | $0.67353 \pm 0.00889$ |
| total | 5 | 0.22 | 0.20 | 5,137 | 3,834 | 2,637 | $0.68779 \pm 0.01183$ |
| 1 | 0 | 0.39 | 0.34 | 185 | 4,443 | 694 | $0.15620 \pm 0.01588$ |
| 1 | 1 | 0.34 | 0.30 | 694 | 5,442 | 1,278 | $0.23484 \pm 0.01138$ |
| 1 | 2 | 0.30 | 0.27 | 1,278 | 2,920 | 910 | $0.31164 \pm 0.01413$ |
| 1 | 3 | 0.27 | 0.24 | 910 | 1,771 | 632 | $0.35686 \pm 0.01882$ |
| 1 | 4 | 0.24 | 0.22 | 632 | 1,063 | 760 | $0.71496 \pm 0.02307$ |
| 1 | 5 | 0.22 | 0.20 | 760 | 498 | 357 | $0.71687 \pm 0.03184$ |
| 2 | 0 | 0.39 | 0.34 | 562 | 13,610 | 1,492 | $0.10963 \pm 0.00780$ |
| 2 | 1 | 0.34 | 0.30 | 1,492 | 11,670 | 2,907 | $0.24910 \pm 0.00819$ |
| 2 | 2 | 0.30 | 0.27 | 2,907 | 6,742 | 2,717 | $0.40300 \pm 0.01010$ |
| 2 | 3 | 0.27 | 0.24 | 2,717 | 5,483 | 2,460 | $0.44866 \pm 0.01125$ |
| 2 | 4 | 0.24 | 0.22 | 2,460 | 3,936 | 2,716 | $0.69004 \pm 0.01221$ |
| 2 | 5 | 0.22 | 0.20 | 2,716 | 1,920 | 1,318 | $0.68646 \pm 0.01662$ |
| 3 | 0 | 0.39 | 0.34 | 28 | 655 | 219 | $0.33435 \pm 0.05720$ |
| 3 | 1 | 0.34 | 0.30 | 219 | 1,654 | 843 | $0.50967 \pm 0.02613$ |
| 3 | 2 | 0.30 | 0.27 | 843 | 1,904 | 1,020 | $0.53571 \pm 0.01881$ |
| 3 | 3 | 0.27 | 0.24 | 1,020 | 1,890 | 1,032 | $0.54603 \pm 0.01874$ |
| 3 | 4 | 0.24 | 0.22 | 1,032 | 1,678 | 1,068 | $0.63647 \pm 0.01924$ |
| 3 | 5 | 0.22 | 0.20 | 1,068 | 792 | 449 | $0.56692 \pm 0.02758$ |
| 4 | 0 | 0.39 | 0.34 | 205 | 5,005 | 648 | $0.12947 \pm 0.01397$ |
| 4 | 1 | 0.34 | 0.30 | 648 | 5,139 | 1,129 | $0.21969 \pm 0.01192$ |
| 4 | 2 | 0.30 | 0.27 | 1,129 | 2,670 | 848 | $0.31760 \pm 0.01489$ |
| 4 | 3 | 0.27 | 0.24 | 848 | 1,813 | 608 | $0.33536 \pm 0.01870$ |
| 4 | 4 | 0.24 | 0.22 | 608 | 950 | 593 | $0.62421 \pm 0.02637$ |
| 4 | 5 | 0.22 | 0.20 | 593 | 624 | 513 | $0.82212 \pm 0.02480$ |

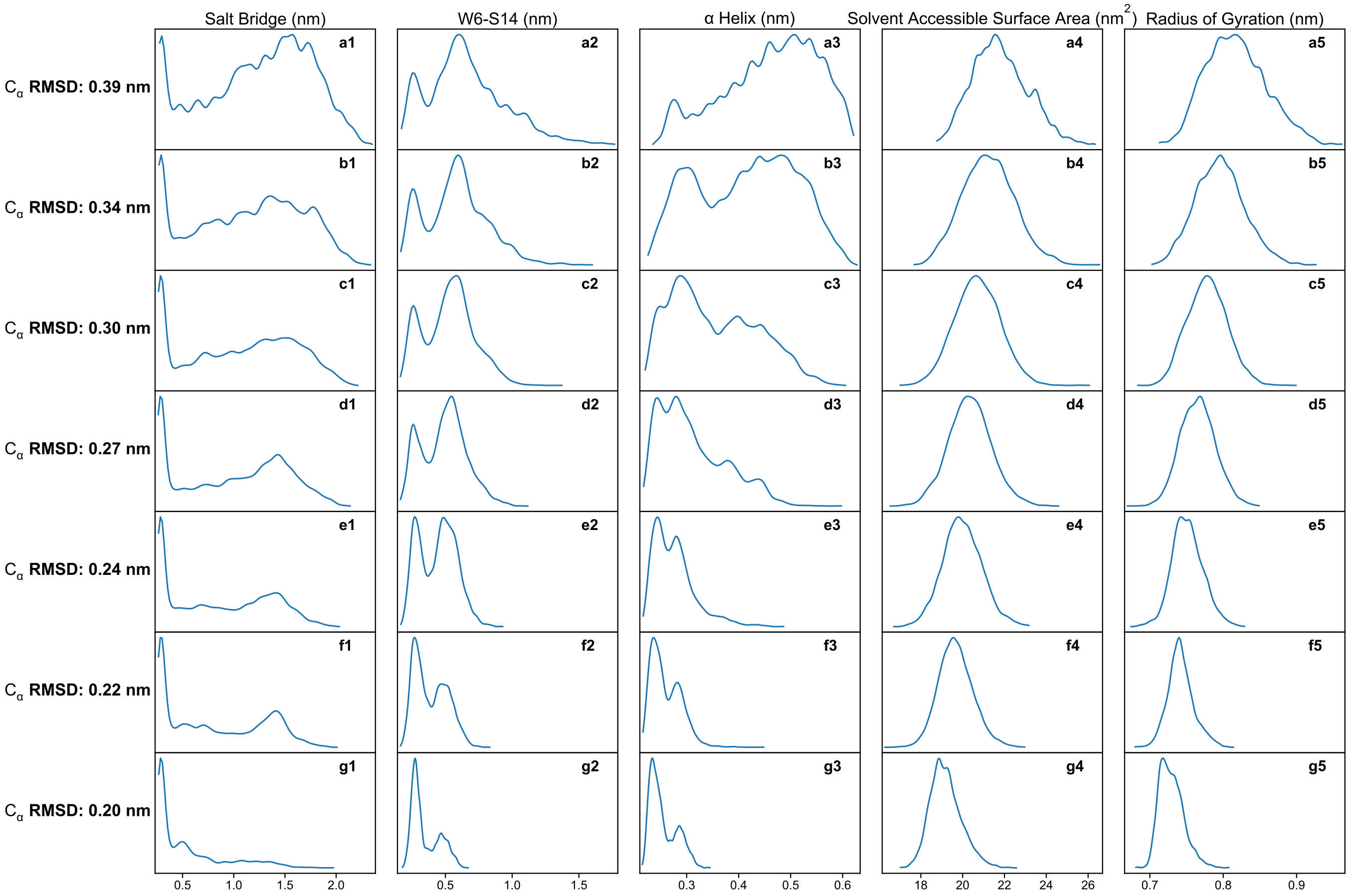


FIG. S2. *Histograms of global and local order parameters at different FFS milestones.*

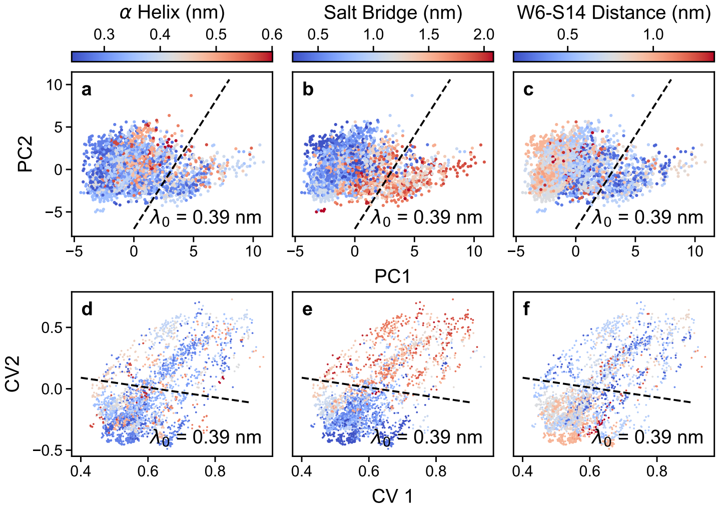


FIG. S3. *Performance of alternative dimensionality reduction approaches.* Scatter plots of the two-dimensional latent spaces obtained from (a-c) PCA and (d-f) a traditional autoencoder colored according to (a,d) $\alpha$-helix RMSD, and (b,e) D9–R16 (salt bridge) distance and (c,f) W6–S14 distance at $\lambda_0 = 0.39\,\text{nm}$. In each panel, an approximate separating line indicating a two-cluster split is overlaid.

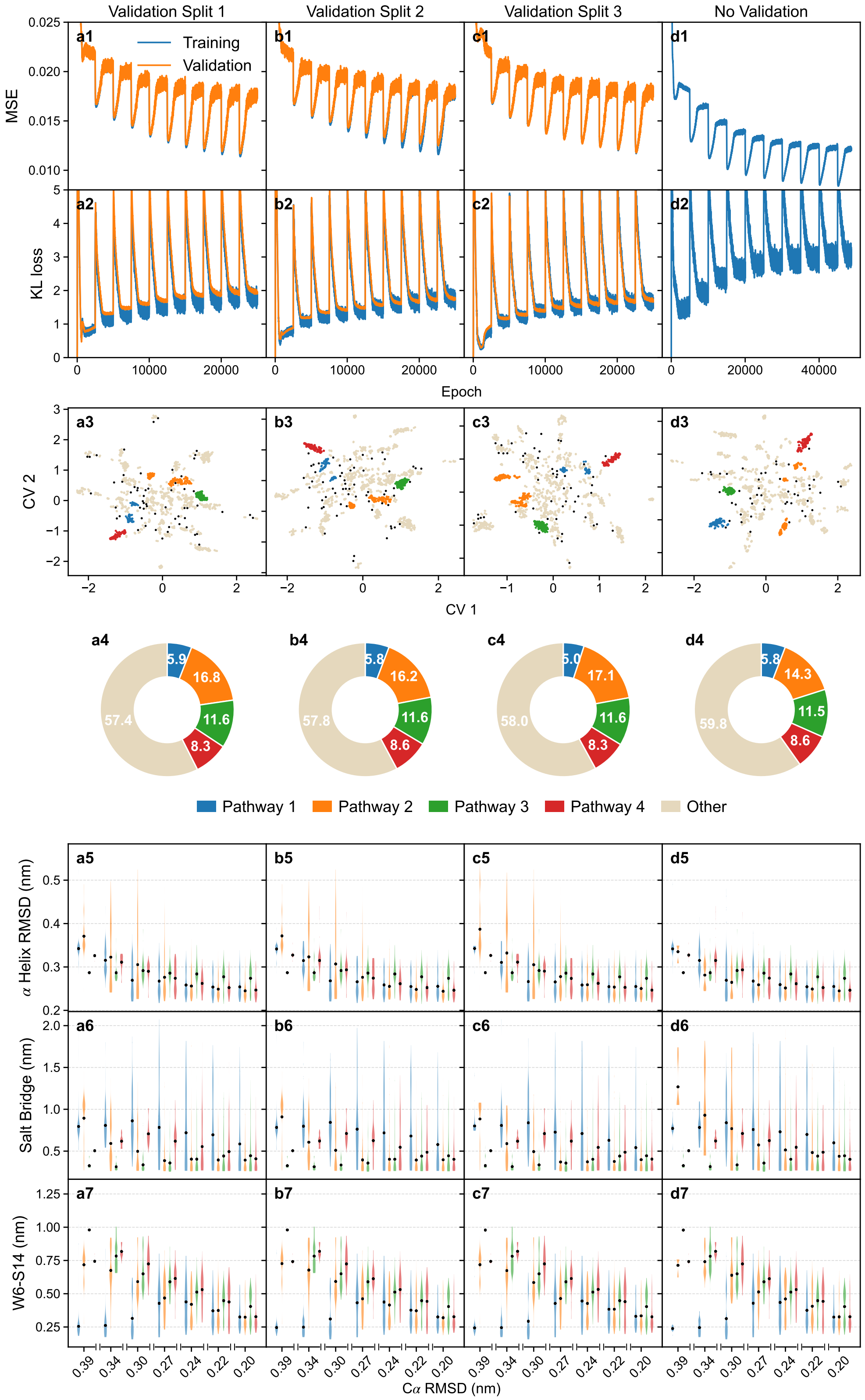


FIG. S4. *Robustness of VAE training.* Evolution of the MSE and KL loss terms with training epoch, together with the resulting latent spaces, fractions of trajectories assigned to major clusters, and violin plots of $\alpha$-helix RMSD, salt-bridge distance, and W6–S14 distance for three independent random splits of the trajectory ensemble into training and test sets. To balance reconstruction fidelity and latent space regularization, the relative weight of the KL divergence term was modulated periodically over ten cycles of 500 training epochs each. This prevents posterior collapse while promoting smooth, informative latent representations. Both MSE and KL losses exhibit a progressive decline across training epochs, with periodic oscillations corresponding to KL ramp-up intervals. These trends reflect progressive improvement in trajectory reconstruction, with sustained nonzero KL values confirming that the latent space remains structured and regularized, enabling low-dimensional projection of folding dynamics. The rightmost column shows training on the full ensemble, yielding indistinguishable loss convergence, latent-space organization, pathway populations, and structural distributions, demonstrating the robustness of the learned representation.

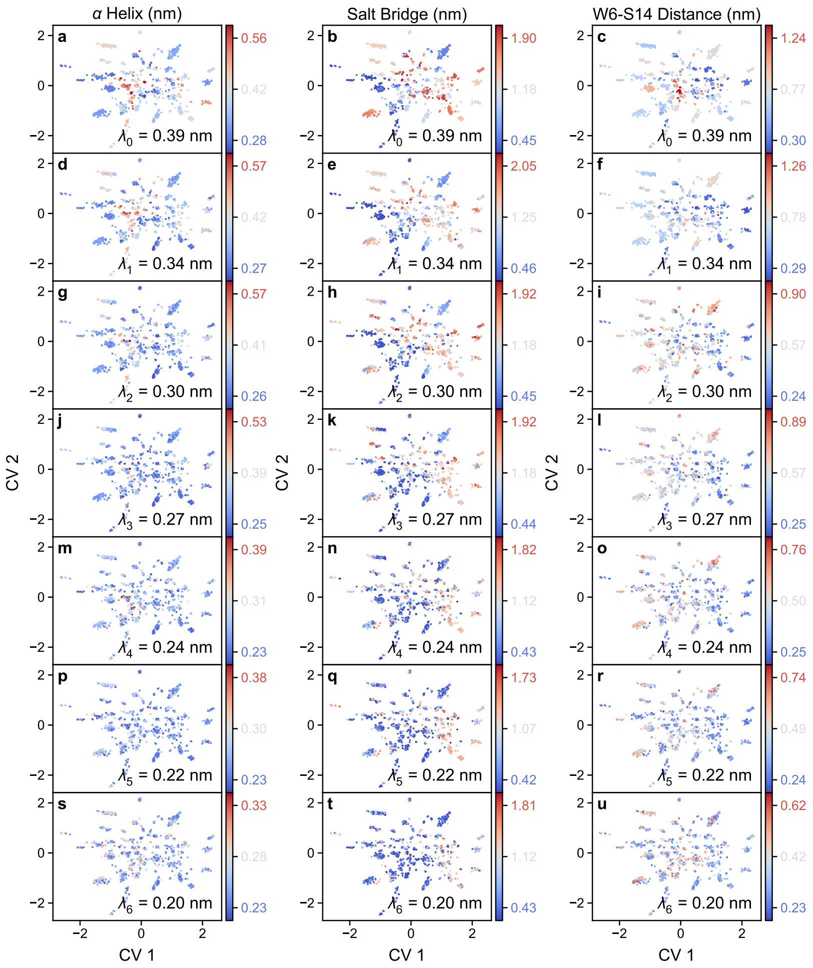


FIG. S5. *Scatter plots of the VAE latent space color-coded based on the values of α-helix RMSD, D9-R16 salt bridge and W6–S14 distances at all FFS milestones.*

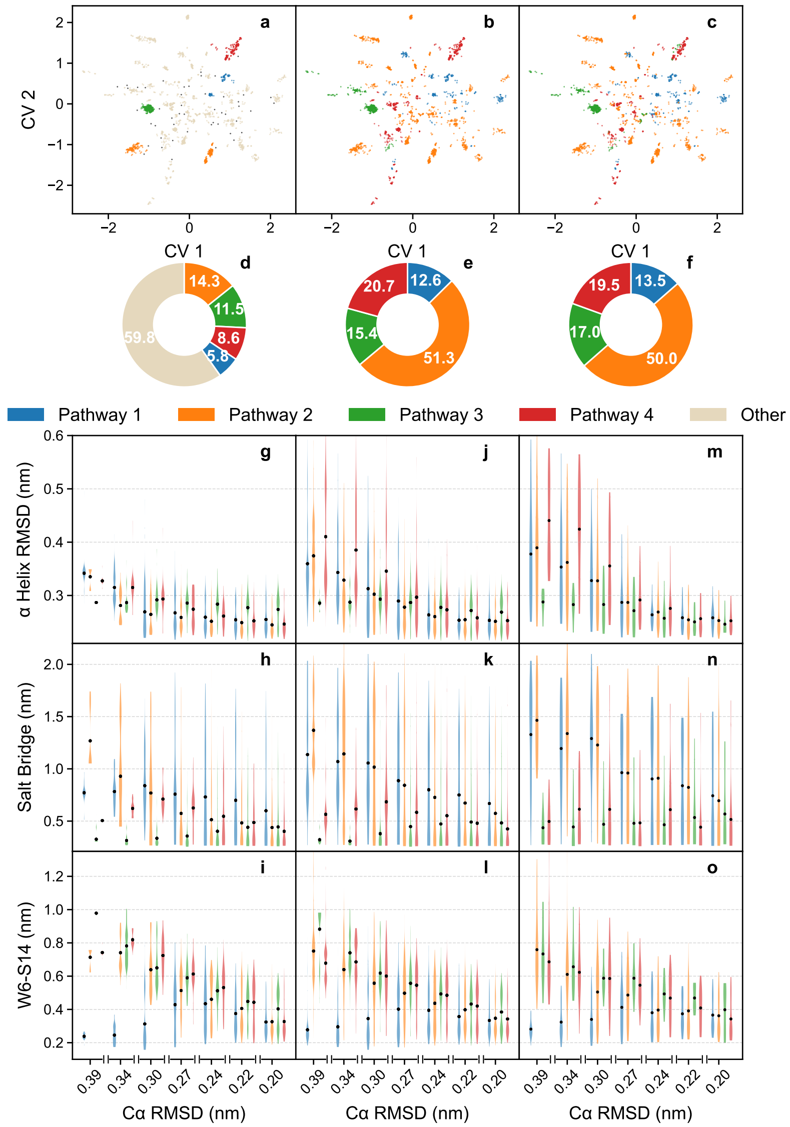


FIG. S6. *Assignment of minor clusters to four major folding pathways.* (a,d,g–i) Density-based clustering in the (a) VAE latent space, together with the (d) fraction of folding trajectories assigned to each major pathway and the corresponding structural distributions shown as violin plots of (g) $\alpha$-helix RMSD, (h) salt-bridge distance, and (i) W6–S14 distance versus $\alpha$-carbon RMSD. Nearly half of all folding events are captured by the four major clusters. (b,e,j–l) Minor clusters assigned to major pathways using mean local order parameters evaluated at $\lambda_0$. (c,f,m–o) Assignment based on trajectory-level order parameters. Although this procedure introduces a small number of latent-space outliers, the pathway populations (e,f) and structural distributions (j–l,m–o) remain largely unchanged.

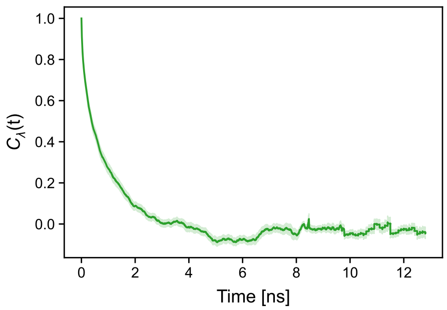


FIG. S7. C$\alpha$ RMSD autocorrelation function computed from basin exploration trajectories. The decorrelation time, corresponding to the decay of $C_\lambda(t)$ to $1/e$ is $\tau = 664^{+64}_{-44}$ ps.
